# Mass univariate testing biases the detection of interaction effects in whole-brain analysis of variance

**DOI:** 10.1101/130773

**Authors:** Robert S. Chavez, Dylan D. Wagner

**Affiliations:** Department of Psychology, The Ohio State University, Columbus, OH, USA

## Abstract

Whole-brain analysis of variance (ANOVA) is a common analytic approach in cognitive neuroscience. Researchers are often interested in exploring whether brain activity reflects to the interaction of two factors. Disordinal interactions — where there is a reversal of the effect of one independent variable at a level of a second independent variable — are common in the literature. It is well established in power-analyses of factorial ANOVAs that certain patterns of interactions, such as disordinal (e.g., cross-over interactions) require less power than others to detect. This fact, combined with the perils of mass univariate testing suggests that testing for interactions in whole-brain ANOVAs, may be biased towards the detection of disordinal interactions. Here, we report on a series of simulated analysis --including whole-brain fMRI data using realistic multi-source noise parameters-- that demonstrate a bias towards the detection of disordinal interactions in mass-univariate contexts. Moreover, results of these simulations indicated that spurious disordinal interactions are found at common thresholds and cluster sizes at the group level. Moreover, simulations based on implanting true ordinal interaction effects can nevertheless appear like crossover effects at realistic levels of signal-to-noise ratio (SNR) when performing mass univariate testing at the whole-brain level, potentially leading to erroneous conclusions when interpreted as is. Simulations of varying sample sizes and SNR levels show that this bias is driven primarily by SNR and larger sample sizes do little to ameliorate this issue. Together, the results of these simulations argue for caution when searching for ordinal interactions in whole-brain ANOVA.

---

Factorial analysis of variance (ANOVA) is the workhorse of statistical methods in psychology and neuroscience. Among the reasons for its widespread use is that it provides a straightforward framework for testing for interactions among levels of different factors (e.g., where the effect of one factor on the dependent variable varies as a function of one or more other factors). For example, within cognitive, social, and affective neuroscience, researchers are often interested in testing whether the effect of a given experimental condition or treatment on brain activity varies as a function of group (e.g., patients vs. healthy participants; young vs. old adults; low vs. high anxiety) or of other conditions (e.g., low vs. high attention, social vs. non-social rewards). Although there are several ways that researchers can apply factorial ANOVA to neuroimaging datasets, perhaps the most common approach is to use ANOVA to identify voxels demonstrating main effects and interactions at the whole-brain level through mass univariate testing. This use of AVNOA is all the more common in instances where researchers have no strong a priori hypothesis about the precise location of these effects or, if they do have hypotheses concerning the spatial location of effects, they may lack the means of identifying independent regions of interest (e.g., localizers, orthogonal contrasts, etc.). In this article, we aim to draw attention to several subtle biases that can arise when testing for interaction effects in whole-brain ANOVA using mass univariate testing. These biases that may lead to spurious interaction effects as well as distortions of true effects thereby obfuscating inferences based on these tests.

Although different areas of scientific inquiry may vary in the precise pattern of interaction that is common to their research area, for instance social psychology often tends to hypothesize disordinal (i.e., crossover) interactions, we believe that a common interaction pattern in social, cognitive and affective neuroscience is the ordinal interaction. Ordinal interactions -- also known as uncrossed, spreading or fan-shaped interactions -- are those in which the size of the effect of one factor varies when under the levels of the other factor. For example, suppose an fMRI study comparing brain activity in young and old adults completing a working memory task at low and high working memory loads. At low working memory loads, it may be expected that there would be minimal differences between younger and older adults, whereas at high loads, these two groups might reasonably be expected to differ. Hypotheses regarding *ordinal* interactions are likely to be the dominant hypothesized interaction type in cases where researchers are comparing groups pre- and post-treatment, drug, or intervention. However, these need not be constrained to between-group studies; there may also be many instances where the effect of a manipulation is expected to differ only under one level of another factor (e.g., the effect of attention on visual search in uncluttered vs. cluttered scenes). Unfortunately, certain features of neuroimaging data analysis may render ordinal interactions far more difficult to detect than disordinal ones and, even in cases where the true effect is ordinal, the resulting statistical parametric map may emphasize voxels that instead show a disordinal interaction due purely to chance and noise..

Why might analyses of neuroimaging data favor the detection of disordinal interactions? To answer that question, we need to discuss the statistical power required to detect different types of interactions. Specifically, the default interaction coefficients in a 2x2 ANOVA represent an omnibus approach to detecting interaction effects that come at the cost of some power to detect specific patterns of interactions, such as ordinal ones (see. Bobko, 1986; Strube & Bobko, 1989). Practically, what this means is that, all things being equal, factorial ANOVAs will be more efficient at detecting disordinal interactions than ordinal ones (Wahlsten, 1991). For example, the required sample size for 90% power to detect an ordinal interaction may be three times larger than that for a comparable disordinal interaction (Wahlsten, 1991). Returning to our working memory example, a plausible ordinal interaction for this study would be one where young adults show no difference in brain activity as a function of working memory load, whereas older adults differ from younger adults only in the high load condition. Another type of ordinal interaction that is perhaps all the more likely, is that both younger and older adults show more brain activity in a given region at high working memory loads -after all it is more difficult-however this effect may be larger for the older adults. This second type of ordinal interaction are even more difficult to detect, and can require nearly four times as many subjects as the previously described ordinal interaction and 10 times as many as a disordinal interaction (Wahlsten, 1991). Thus, in general, the traditional interaction coefficients in an ANOVA tend to favor disordinal interactions, requiring smaller sample sizes to detect these with reasonable power. This bias to detect certain types of interactions over others presents a problem in neuroimaging research where multiple comparisons are often an issue. That is to say, when searching for the most significant interaction effects across thousands of statistical tests, a disproportionate number of disordinal interactions are likely to be significant, regardless of the true underlying relationship of the phenomenon. Even in cases where the data represent only noise, false positive voxels are expected to favor spurious disordinal interactions over ordinal ones.

This particular issue seems to have l been overlooked in the literature. It is also unclear to what degree this bias may be compounded by the various sources of variability that contribute to the inherent noisiness of the BOLD signal in fMRI data, including: head movement, physiological noise, scanner noise, and individual variability in the BOLD responses among subjects. In this article, we demonstrate the scope of this phenomenon in simulated datasets. First, to illustrate the general phenomenon, we show that disordinal interactions are strongly overrepresented among the top significant tests using a simulated dataset of 10,000 “voxels” consisting only of pure Gaussian noise. Next, we inject a known true ordinal interaction into the simulated datasets and re-run the analyses using different levels of signal to noise ratio (SNR) and sample sizes in order to explore how mass univariate testing can distort ordinal interactions towards a disordinal interaction pattern as a function of the different noise parameters and sample sizes. Finally, to demonstrate that these biases obtain in realistic fMRI data and can survive cluster correction, we use an open source package for generating simulated fMRI data (neuRosim; Welvaert, Durnez, Moerkerke, Verdoolaege, & Rosseel, 2011) with realistic noise properties and injected an ordinal interaction into several regions of interest (ROI). These simulated datasets were then analyzed in a common neuroimaging analysis package (FSL) and submitted to a 2^nd^ level group analysis examining clusters showing significant interactions.

## Methods

In this article, we employ two sets of simulated ANOVA analyses. The first set were run to demonstrate general issues with detecting ordinal interactions during mass univariate testing, without respect to the influence of the fMRI signal or the spatial characteristics of the brain. The second set used simulated fMRI datasets to explore the effects of spatial clustering on both real and spurious interactions in realistic fMRI data after preprocessing and statistical modeling using standard fMRI analysis tools.

### Simulations in pure noise data

Before highlighting how issues with detecting ordinal interactions may influence whole-brain fMRI data, we first sought to demonstrate the presence of this phenomenon in a simpler framework that mimics the mass univariate testing approach common in fMRI. To accomplish this, we ran a series of ANOVA simulations in both pure noise data and in data where a known true ordinal interaction effect was implanted in the data. All simulations were conducted in the R statistical language (R Development Core Team, 2011).

First, we sought to show that spurious disordinal interactions are disproportionately detected relative to other types interaction effects when sorting by the most significant effects in pure noise -an approach that mimics mass univariate testing in fMRI where typically only the voxels with the smallest p-values survive correction for multiple comparisons. In total, 10,000 simulated ANOVA tests were conducted on pure noise data. Within each test, the dependent measure was generated by randomly sampling a from a uniform distribution *U*(*0*,*10*) in order to produce a pure noise dataset^1^. These values were assigned to one of four condition cells with an N = 40 per cell. We then conducted a standard 2×2 ANOVA for each simulation and extracted the statistics for each interaction term in the model.

### Simulations in data with a known true ordinal interaction

In addition, we also ran a series of simulations that included the presence of a known ordinal interaction effect at multiple sample sizes and signal-to-noise (SNR) levels. The simulated effect compared two hypothetical conditions at two levels where there was no difference between conditions at the first level and a one-unit difference between the two conditions at the second level (see: Figure 1). Simulated data were randomly sampled from a Gaussian distribution with μ = 0 for each cell except the one driving the interaction effect which was sampled at μ = 1, and standard deviations of the Gaussian distributions were set to either σ = 8, σ = 4, σ = 2, or σ = 1. SNR was defined as the ratio of the difference between the means of the Gaussian distributions and their standard deviation set at each simulation. These simulations were repeated at sample sizes of 20, 40, 60, 80, and 100. A set of 1000 simulations were run at each of four levels of SNR and five sample sizes for a total of 20,000 ANOVA across all tests.

**Figure 1.**
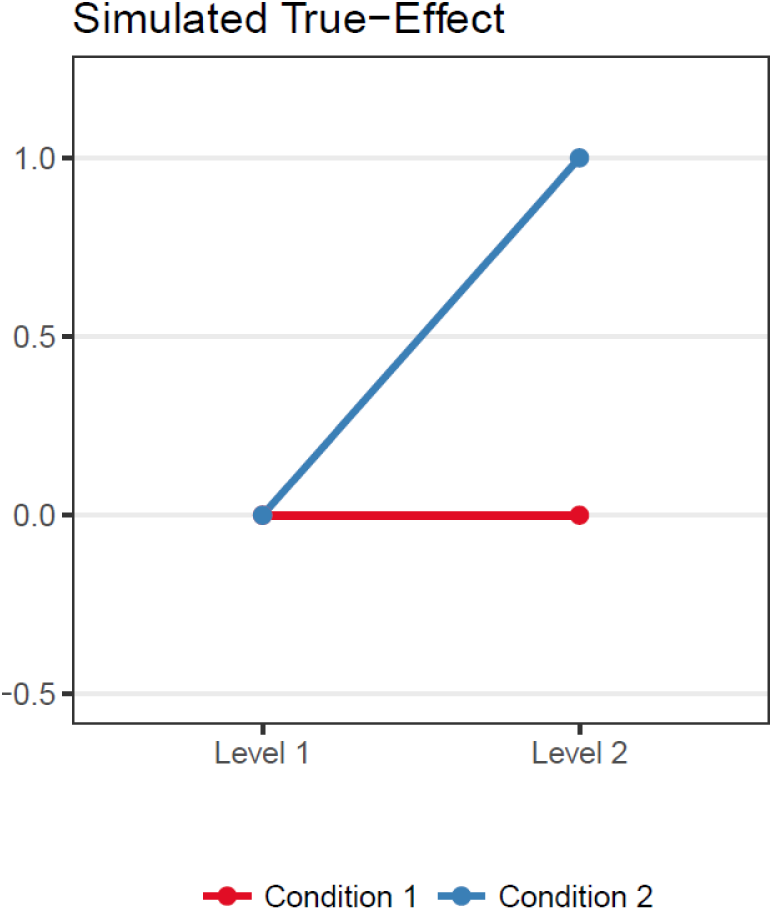
The shape of the interaction effect used in the simulations in which a true-effect was present.

### Simulation and analysis of realistic fMRI data

Simulations of fMRI data were conducted using the R package neuRosim (Welvaert et al., 2011). The simulated fMRI scans consisted of 216 total volumes with a 2 second TR. Voxels size was set to 3mm isotropic in a total brain volume spanning 60 × 70 × 55 voxels. Simulated scanner noise was composed of a mixture of noise sources used in the original neuRosim publication (Welvaert et al., 2011, 2011), which included 30% Rician system noise (Gudbjartsson & Patz, 1995), 30% temporal noise, 1% low-frequency drift, 9% physiological noise, 1% residual head movement artifacts, 2% spatially correlated noise, and with an overall SNR = .5. We simulated a 2×2 within-subjects ANOVA analysis in an event-related design consisting of two levels each of two experimental conditions with 30 events per condition per level in a total of N = 100 simulated subjects. The shape of the fMRI simulated interaction effect is the same as the one specified in the non-fMRI simulations and illustrated in Figure 1. Specifically, each event was simulated to invoke a mean BOLD response of μ = 1 unit above baseline except for the cell driving the interaction effect which had a mean BOLD response of μ = 2 units above baseline. Inter-subject variability in these responses was simulated by sampling each of these BOLD responses from a Gaussian distribution with standard deviation of σ = 1 and centered on their respective means, note that this variability models the size of the effect and not individual variability in the shape and duration of the BOLD response which was set to the same parameters for all subjects. These effects were implanted in five spherical regions of interest (ROIs) each with a diameter of 3 voxels. The remaining voxels outside of these ROIs consisted of only of simulated noise.

These simulated fMRI datasets were preprocessed and analyzed using FSL. Preprocessing consisted of spatial smoothing using a Gaussian kernel of FWHM 6 mm^3^; mean-based intensity normalization of all volumes by the same factor; and high-pass temporal filtering (Gaussian-weighted least squares running line smoother with sigma = 90s). Time-series statistical analysis was carried out using a regularized autocorrelation model with prewhitening. Because all simulated data were generated in a common coordinate space, motion correction and registration to a standard stereotaxic space were not applicable. First-level parameter estimate maps were calculated using the general linear model by contrasting the main effects and their interaction from each factor in the ANOVA. Group-level statistical analysis was conducted using non-parametric permutation tests with FSL’s Randomise (Winkler, Ridgway, Webster, Smith, & Nichols, 2014), using an initial cluster-extent forming threshold of *p* <.001 (uncorrected). Cluster-level thresholding was set to *p* <.05 corrected for family-wise error rate. These parameters were chosen to mimic preprocessing and analysis parameters that are common in the functional neuroimaging literature.

## Results

### Interaction effects in pure noise data

A 2×2 ANOVA was calculated on 10,000 simulated datasets composed of pure Gaussian noise, mimicking the mass univariate testing approach common to functional neuroimaging research. The results of these analyses were sorted by individual simulations with the lowest *p*-values for the interaction term. Of the 10,000 simulations conducted, 565 tests produced an interaction effect of *p* <.05, yielding a family-wise error rate of 5.65%. In order to identify the shape of these 565 significant interactions, we opted to define disordinal interactions as those in which the mean of each level of one condition was beyond one standard error of the other condition. We could have tested the simple effects, however because it is possible to have a disordinal interaction in the absence of significant simple effects, we chose this approach as being the most straightforward. Based on this definition of disordinal interactions, we found that 387 of the 565 (69%) of the significant interaction effects presented a disordinal interaction pattern. In addition, we also examined the results for the fifty most significant interactions (Figure 2). Although these analyses were modeled using only randomly generated noise with no true signal, it is clear that disordinal patterns dominate among significant interaction effects, especially when these results are filtered by the significance of the interaction term.

**Figure 2.**
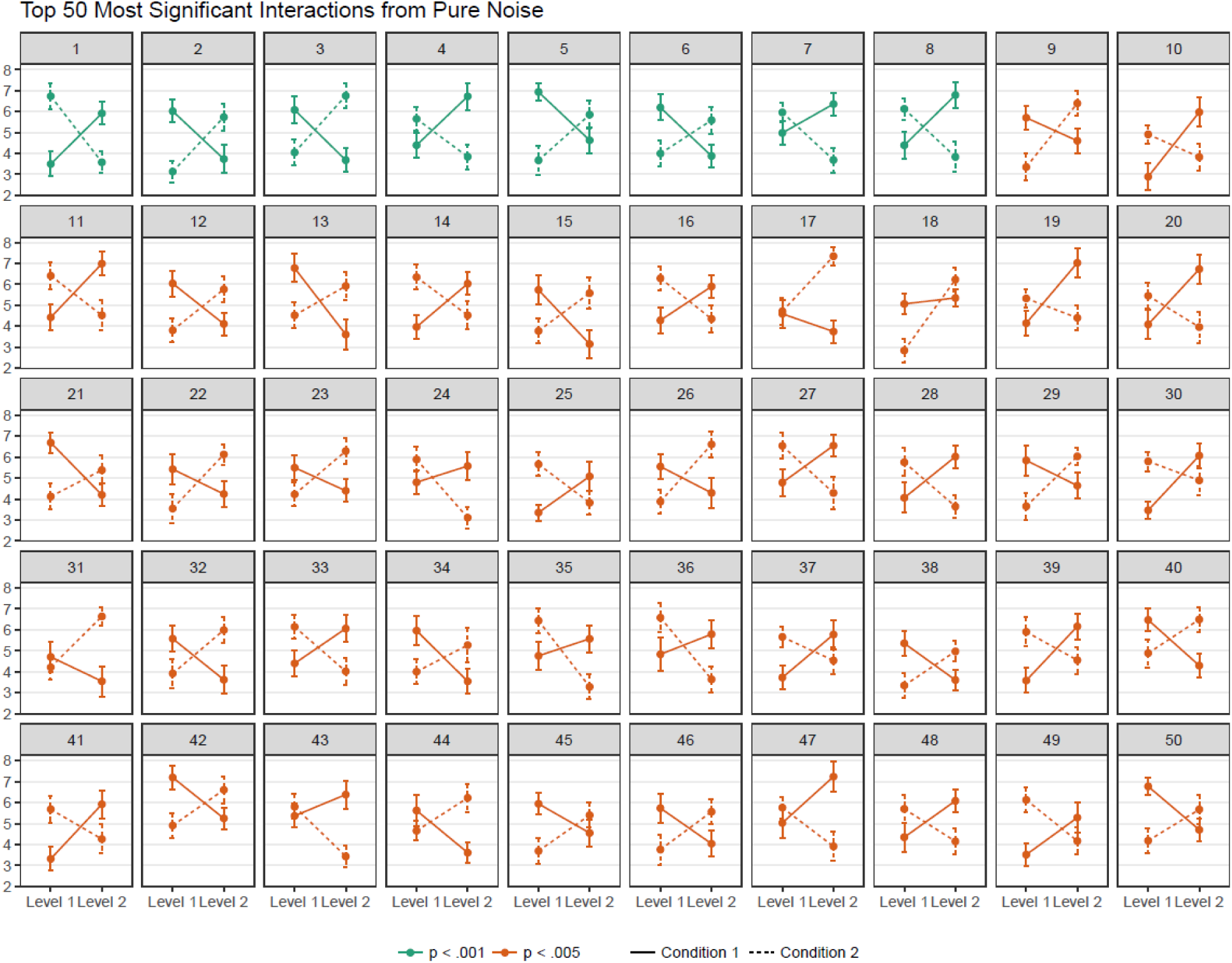
Results of the top fifty most significant interaction effects from the pure noise data. Compared to other kinds of interactions, disordinal interactions are disproportionately represented when filtering by significance.

### Interactions effects in data with a known true ordinal interaction

The results from the simulations in which a true-effect is present were calculated in the same manner as the pure noise data described above with the exception that we also tested the effect of sample size and SNR levels on the amount and shape of the significant interaction effects. Among each of the interaction effects with a *p* <.05, we calculated the proportion of disordinal interactions for each SNR and sample size combination. Results from these calculations are shown in Table 1 for two common cluster forming thresholds in neuroimaging (i.e., p<0.001 and p<0.005). In general, lower SNR had led to a greater number of disordinal interactions tests, whereas sample size had less of an influence. In order to emulate the effects of significance filtering on these data, the fifty most significant interaction effects within each of SNR/sample size combination were then averaged together to show the shapes of the interaction effects when aggregated across the most significant results. These results are displayed in Figure 3 and indicate that as SNR decreases, significant interaction effects are more likely to resemble a disordinal interaction when results are filtered based on significance level, even in the presence of a true-effect as in these simulations. Moreover, these results also show that increases in the sample size of the data appear to do little to mitigate this problem. At lower SNR values, large sample sizes narrow the confidence interval around individual condition means but do not eliminate the distortion of true ordinal interaction towards a disordinal interaction pattern.

**Table 1.** Percentage of disordinal interaction effects among interactions in simulated datasets at varying SNR and sample sizes when a true ordinal interaction was present.

| SNR | N = 20 |  | N = 40 |  | N = 60 |  | N = 80 |  | N = 100 |  |
| --- | --- | --- | --- | --- | --- | --- | --- | --- | --- | --- |
|  | p<0.001, p<0.005 |  | p<0.001, p<0.005 |  | p<0.001, p<0.005 |  | p<0.001, p<0.005 |  | p<0.001, p<0.005 |  |
| 0.125 | N/A | 75.05 | 100 | 100 | 100 | 100 | 100 | 81.8 | 100 | 85.7 |
| 0.25 | 100 | 86.7 | 100 | 96.0 | 91.7 | 89.7 | 75 | 75.0 | 78.8 | 73.8 |
| 0.5 | 100 | 92.8 | 81 | 71.8 | 72.4 | 60.7 | 65.7 | 53.8 | 58.7 | 47.9 |
| 1 | 67.4 | 55.6 | 48.1 | 40.1 | 31.2 | 25.6 | 24.4 | 22.3 | 22.9 | 22.0 |

*Note:* For each cell percentages are displayed for common fMRI thresholds of $p<0.001$ (left) and $p<0.005$ (right). N/A refers to a simulated dataset where no p-value passed the threshold of $p<0.001$ .

**Figure 3.**
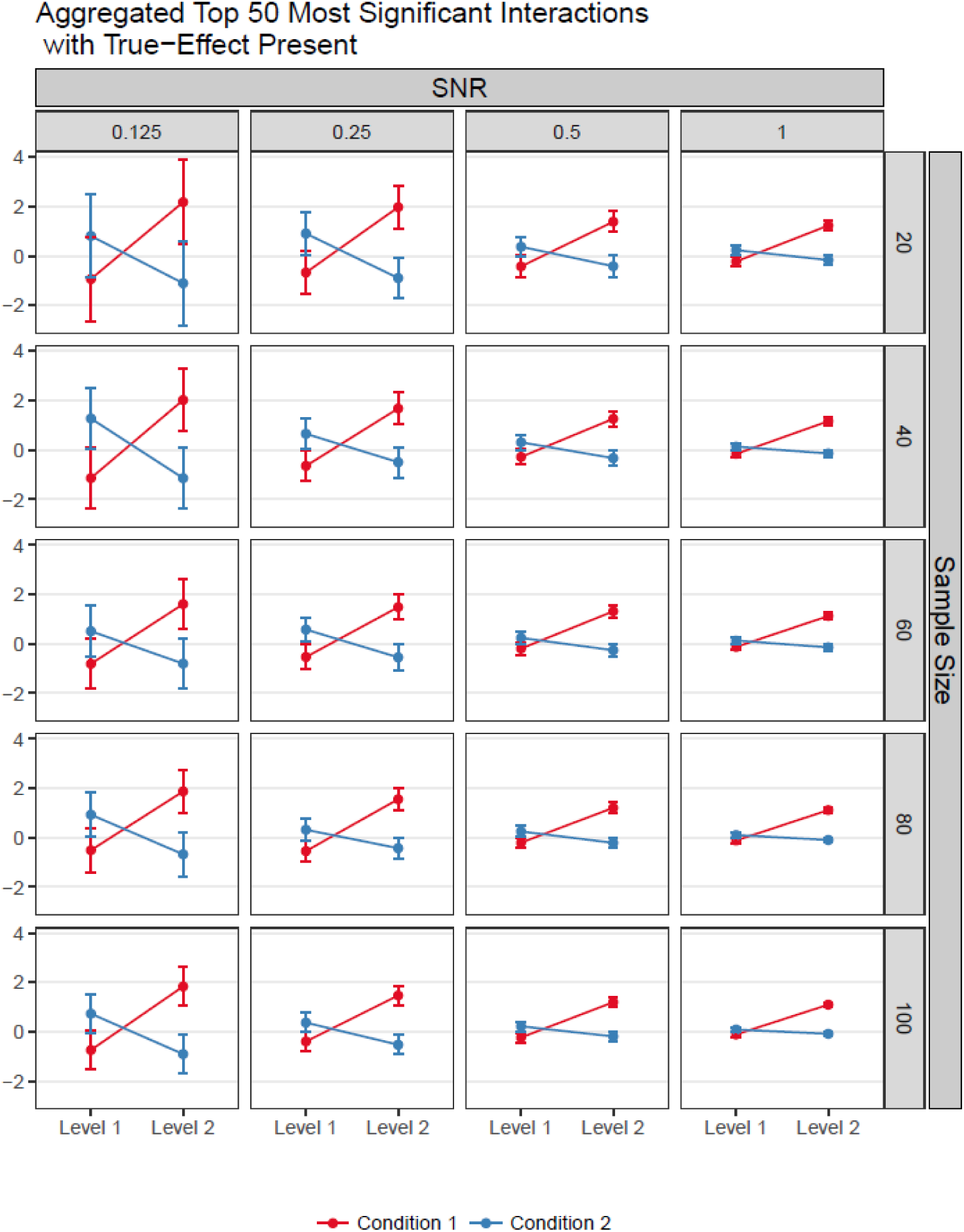
Results from the simulations containing a true known ordinal interaction after significance filtering. Within each sample size and SNR level, the average effects and standard errors for the top fifty most significant interactions are displayed. As SNR levels decrease, significant interactions are more likely to resemble spurious disordinal interactions. Sample size appears to do little to mitigate this bias.

### Interaction effects in stimulated fMRI data with known true ordinal interaction effect

In this analysis, we generated one hundred separate fMRI datasets using a realistic mode of fMRI noise (as implemented in neuRosim, see methods). For each of these simulated subjects, we generated a dataset for and event-related design with 2 factors each with two levels (i.e., four conditions total), and containing thirty separate events. Within these datasets, we simulated a true ordinal interaction effect and injected it into five spherical ROIs (diameter: 3 voxels) and seeded throughout the brain. To best approximate the analysis of real fMRI data, these simulated datasets were then made to undergo a standard preprocessing and 1^st^ and 2^nd^ level analysis in the FSL software package (see methods). In this way, the results of the analyses on these simulated datasets are performed using the same software, preprocessing steps, and statistical modeling approaches as in fMRI analyses with real data. This analysis therefore allows us to answer the question of whether the biases towards disordinal interactions revealed in the non-fMRI simulations above would obtain in a standard fMRI analysis and whether they would spatially cluster in sufficient number to survive cluster correction.

The results of a group level analyses of the 100 simulated subjects produced seven significant clusters in which a 2×2 significant interaction effect was identified using non-parametric permutation tests (See Figure 4). Of these clusters, five of them reflected the location of the implanted true-effects and two clusters were false positives. Parameter estimates for each experimental condition averaged within each cluster are plotted in Figure 5. Within the five true positive clusters, the interaction effects more closely resemble the true-effect implanted within the data, although with a clear tendency to take on a more disordinal or crossover shape. That said, in no case was the difference between the two conditions at the 1^st^ level of each factor significant. Conversely, the two false positive clusters show a clear disordinal interaction shape. These findings are consistent with the results from the non-fMRI simulations and show that spurious disordinal interactions can survive cluster correction, even when using non-parametric cluster correction (e.g., Eklund, Nichols, & Knutsson, 2016) and a comparatively large sample size.

**Figure 4.**
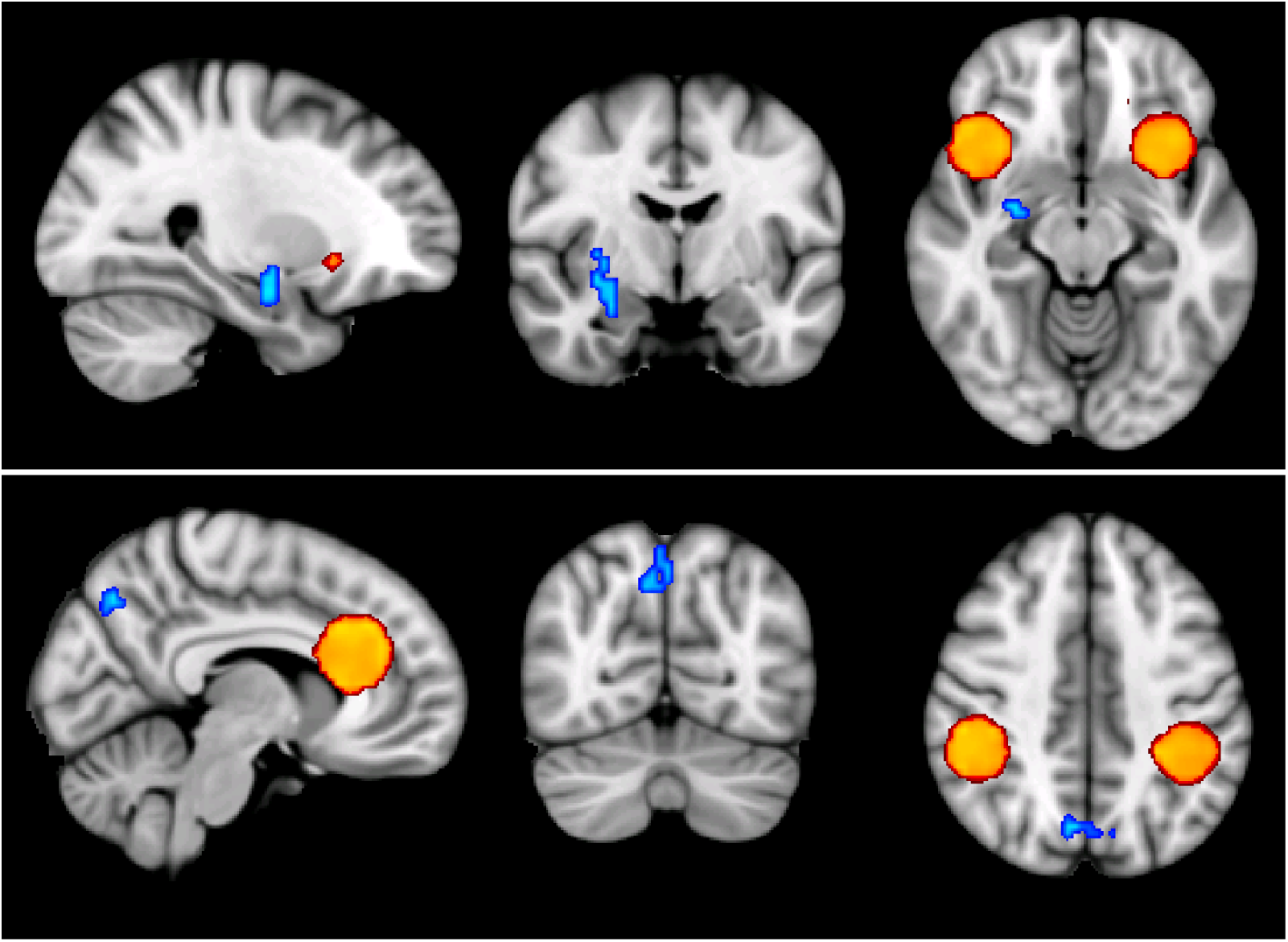
Significant clusters identified from the whole-brain analysis of the simulated fMRI data overlaid on a template brain. The five true positive clusters were identified and are displayed in warm colors. Two false positive clusters survived multiple comparison correction and are displayed in cool colors. (Note: Because these data were generated in simulated space, locations of all clusters are arbitrary with respect to the actual neuroanatomy.)

**Figure 5.**
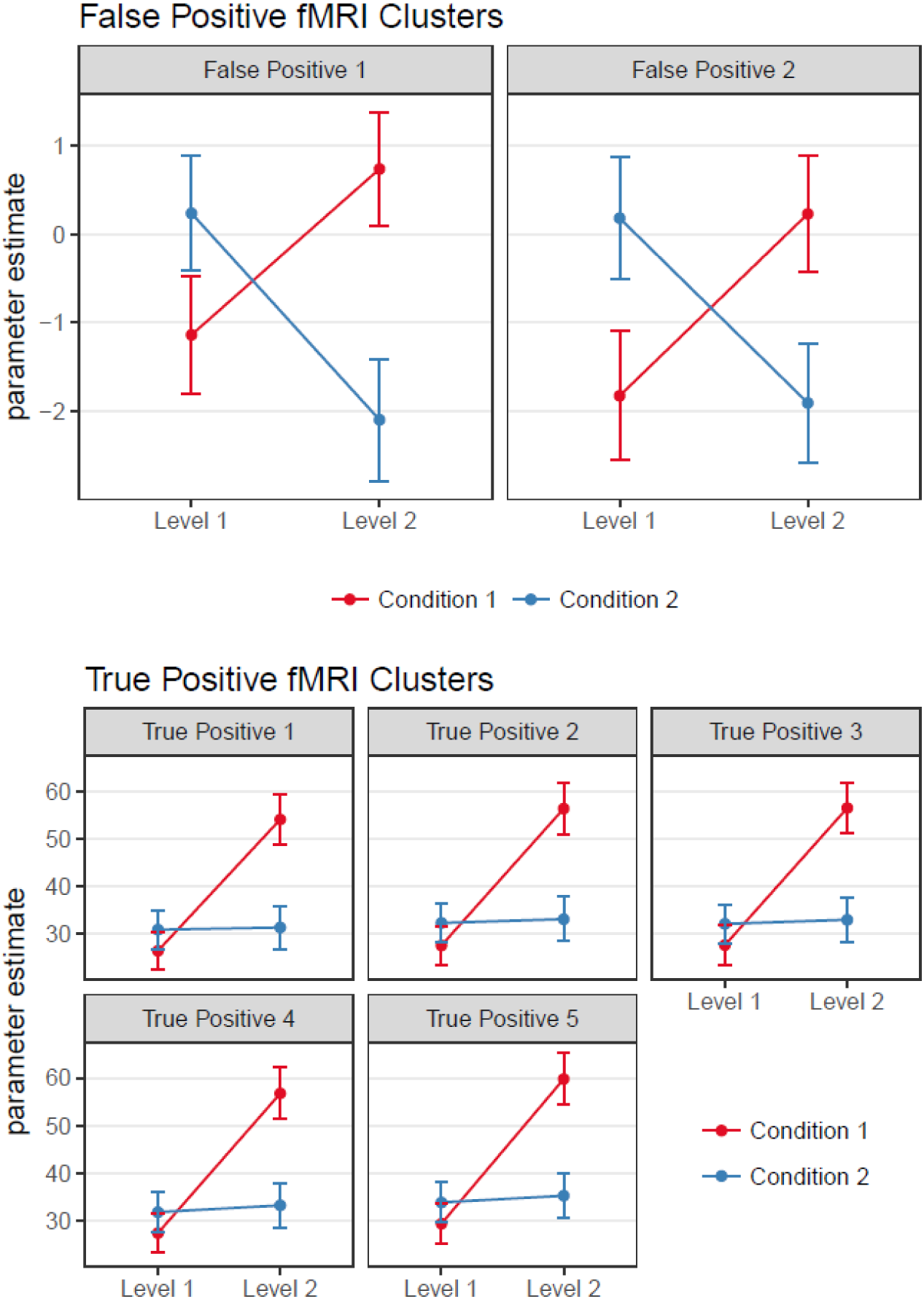
Parameter estimate extracted from the seven significant clusters identified in the in the whole brain analysis. The two false positive clusters show a marked disordinal interaction shape, whereas the five true positive clusters more closely resemble the true effect.

## Discussion

The results of the current study demonstrate that fMRI whole-brain ANOVA analyses are biased towards detecting disordinal interaction effects. In a simulated set of non-fMRI data consisting of pure noise, disordinal interactions dominated the proportion of false positives detected. When simulating data with a known true effect, significance filtering leads to true ordinal interactions beginning to appear like disordinal interactions in measurements with low SNR. Finally, when simulating fMRI data using realistic multi-source noise patterns, despite having a large sample of 100 simulated subjects, spurious disordinal interactions were found using widely-accepted cluster forming thresholds and non-parametric permutation tests. Together, these results raise several concerns about the use of traditional whole-brain analysis procedures when testing for interaction effects is of primary interest.

Although the specific issues with whole-brain ANOVA raised here have not, to our knowledge, been described previously, some of the problems underlying these issue are well known. The disordinal interaction bias shown here appears to arise at the intersection of two well-established issues in both neuroimaging and ANOVA methods more generally. First, it is well-established that whole-brain searches identifying regions showing the lowest *p*-values tend to lead to inflated effect size estimates when they are not calculated in an independent dataset (Kriegeskorte, Simmons, Bellgowan, & Baker, 2009; Vul, Harris, Winkielman, & Pashler, 2009). Second, as discussed above, disordinal ANOVA interactions generally require smaller sample sizes to detect (Wahlsten, 1991) making them more likely to be disproportionally represented in mass univariate tests. In the case of whole-brain ANOVA, these two issues come together to produce a situation in which a whole-brain search identifies areas with inflated interaction effect sizes which are, in turn, more likely to resemble disordinal interaction shapes than other type of relationships, due to the simple fact that these are easier to detect using standard ANOVA interaction coefficients. Moreover, our results also show that these effects seem to be primarily driven by SNR and that larger sample sizes do little to ameliorate this issue. Given the noisy nature of the BOLD response, and of fMRI data more generally, it seems unlikely that subtler ordinal interaction effects would be able to be accurately identified using a whole-brain search procedure, even if they truly exist. Some additional issues that may contribute to a greater bias than estimated here, are the fact that our simulated datasets do not simulate anatomical variability. The location of implanted ordinal interactions in our simulated fMRI subjects was at the same location for all subjects. In real-world fMRI datasets, functional regions are known to vary across subjects both as a result of individual differences in cortical location of functional regions, as well as with inaccuracies related to aligning different subjects’ brains to a common stereotaxic space. For example, if all subjects showed a true ordinal interaction effect in the DLPFC, variance in the precise spatial location of the DLPFC across subjects would give a foothold to noise and chance and would, we argue, be more likely to push the shape of any discovered interaction effect towards that of a disordinal one. Moreover, the current simulations assumed an identical BOLD response across all participants and does not capture true variance in the shape or timing of the underlying hemodynamic response that exists among individuals.

In this article, we focused exclusively on ordinal interactions where there is assumed to be no difference between one level of each factor while the other levels differ between factors. Put more concretely, imagine we had an experiment whereby two groups completed a task before and after a manipulation that in one group was easy, and in another group required the sustained exertion of self-control. At time 1 we assume that random assignment would lead to no differences between group in terms of brain activity, whereas at time 2, the two groups may differ with the self-control exertion group showing more (or less) brain activity than controls. In this example, we may presume that the control participants showed no real change in brain activity as the manipulation was not designed to impact them. In this case the ordinal interaction would take the shape shown in Figure 1. However, in many real-world examples, it is much more plausible to hypothesize that both groups will be impacted by the manipulation but by different amounts. These types of ordinal interactions have been shown to be much more difficult to detect and require a much larger sample size than the ordinal interaction we specified in Figure 1 (Wahlsten, 1991). Based on the simulations and analyses in this article, we can only but conclude that attempting to detect this latter type of ordinal interactions would require incredibly large sample sizes to have sufficient power and may still fall prey both spurious disordinal interactions and a tendency for true ordinal interactions to be distorted towards a disordinal pattern.

Can these statistical biases be avoided? One possibility that may be considered is the use of planned contrasts (e.g., 1 1 1 -3) designed specifically to detect an ordinal pattern of results. However, we caution against this method as prior studies investigating this approach in behavioral experiments have argued that they tend to have an inflated family wise error rate (Strube & Bobko, 1989) due in part to a the fact that they can be “fooled” by a range of interaction types and may produce a significant interaction in cases that are clearly not ordinal in the way hypothesized by the planned contrast (see Petty, Fabrigar, Wegener, & Priester, 1996). In the context of mass univariate testing, these two facts together would be exacerbated as any method with a high false positive rate and noted lack of specificity would be likely to produce a large amount of spurious interactions when used with functional neuroimaging data.

It is our opinion that the best means of avoiding this phenomenon is to restrict the search space and not perform mass univariate testing. Restricting the analyses to several a priori ROIs, be they from localizers, anatomy or orthogonal contrasts, presents the best way of skirting the tendency in mass univariate testing to detect spurious disordinal interactions and distortions of true ordinal ones. In many ways, the effects discussed in this paper are a special case of non-independent analyses (e.g. Kriegeskorte et al., 2009) whereby effect sizes tend to be inflated when performing non-independent ROI analyses. That said, we believe the particular bias towards disordinal interactions in factorial ANOVA designs has not been readily appreciated as indicated by multiple studies detecting and interpreting disordinal interactions in cases where theory or random assignment might have predicted an ordinal interaction pattern. Thus, for researchers who are accustomed to testing their hypotheses using complex interaction models in behavioral data, testing similar models using fMRI presents additional challenges when performing mass univariate testing, challenges that may lead to mistaken inferences of between condition differences at the level of one factor where none exist.

1 Note: We used a Uniform distribution here as a proof of concept that this issue can arise from uniform noise and is not simply an issue of sampling from the Gaussian distribution. Indeed, sampling from a Gaussian distribution does not affect the underlying issue, and we suspect this would be true for other distributions as well.

## Additional Information

Code to perform the non-fMRI simulation analyses and generate the simulated fMRI datasets have been made available here: https://github.com/robchavez/ANOVA_simulations.

## References

Bobko, P. (1986). A solution to some dilemmas when testing hypotheses about ordinal interactions. Journal of Applied Psychology, 71(2), 323–326. https://doi.org/10.1037/0021-9010.71.2.323

Eklund, A., Nichols, T. E., & Knutsson, H. (2016). Cluster failure: Why fMRI inferences for spatial extent have inflated false-positive rates. Proceedings of the National Academy of Sciences, 113(28), 7900–7905. https://doi.org/10.1073/pnas.1602413113

Gudbjartsson, H., & Patz, S. (1995). The rician distribution of noisy mri data. Magnetic Resonance in Medicine, 34(6), 910–914. https://doi.org/10.1002/mrm.1910340618

Kriegeskorte, N., Simmons, W. K., Bellgowan, P. S. F., & Baker, C. I. (2009). Circular analysis in systems neuroscience: the dangers of double dipping. Nature Neuroscience, 12(5), 535–540. https://doi.org/10.1038/nn.2303

Petty, R. E., Fabrigar, L. R., Wegener, D. T., & Priester, J. R. (1996). Understanding Data When Interactions Are Present or Hypothesized. Psychological Science, 7(4), 247–252. https://doi.org/10.1111/j.1467-9280.1996.tb00368.x

R Development Core Team, R. (2011). R: A Language and Environment for Statistical Computing. R Foundation for Statistical Computing (Vol. 1). https://doi.org/10.1007/978-3-540-74686-7

Strube, M. J., & Bobko, P. (1989). Testing hypotheses about ordinal interactions: Simulations and further comments. Journal of Applied Psychology, 74(2), 247–252. https://doi.org/10.1037/0021-9010.74.2.247

Vul, E., Harris, C., Winkielman, P., & Pashler, H. (2009). Puzzlingly High Correlations in fMRI Studies of Emotion, Personality, and Social Cognition. Perspectives on Psychological Science, 4(3), 274–290. https://doi.org/10.1111/j.1745-6924.2009.01125.x

Wahlsten, D. (1991). Sample size to detect a planned contrast and a one degree-of-freedom interaction effect. Psychological Bulletin, 110(3), 587–595. https://doi.org/10.1037/0033-2909.110.3.587

Welvaert, M., Durnez, J., Moerkerke, B., Verdoolaege, G., & Rosseel, Y. (2011). neuRosim: An R package for generating fMRI data. Journal of Statistical Software, 44(10), 1–18.

Winkler, A. M., Ridgway, G. R., Webster, M. A., Smith, S. M., & Nichols, T. E. (2014). Permutation inference for the general linear model. NeuroImage, 92, 381–397. https://doi.org/10.1016/j.neuroimage.2014.01.060

